# Contrasting patterns of seascape genetics in *Acropora cf. tenuis* and their symbiotic algae

**DOI:** 10.64898/2026.04.07.716991

**Authors:** Jaelyn T. Bos, Lisa C. McManus, Rachel Ravago-Gotanco, Malin L. Pinsky

**Affiliations:** Department of Ecology & Evolutionary Biology, University of California Santa Cruz, Santa Cruz, California, USA.; University of Hawaiʻi at Mānoa School of Ocean and Earth Science and Technology, Hawaiʻi Institute of Marine Biology, Kāne‘ohe, Hawaiʻi, USA.; The Marine Science Institute, University of the Philippines, Diliman, Quezon City, Philippines.

## Abstract

Theory suggests that pairs of mutualist species will often co-disperse and share the same dispersal patterns, though the extent to which this happens remains unclear. Photosynthetic corals constitute an example of this dynamic, as they rely on algal symbionts to meet their energetic needs, yet many acquire their symbionts environmentally after their larval dispersal phase. Consequently, corals and their symbionts may exhibit similar or contrasting patterns of genetic variation across the seascape, with implications for their evolutionary and ecological processes. Here, we densely sample corals of the key reef-building taxon *Acropora cf. tenuis* and their algal symbionts across a reefscape in the central Philippines to examine genetic variation across space. Four distinct coral taxa show genetic evidence of long distance dispersal, including weak or absent isolation by distance signals and parent-offspring pairs at widely spaced sites. These coral taxa all host a single group of algal symbionts from the genus *Cladocopium*, which shows landscape genetic structure independent from its coral hosts. In fact, *Cladocopium* genetics vary with both latitude and depth, potentially indicating genome-wide local adaptation at a finer spatial scale than that seen in their hosts. Genetic variation at markedly different spatial scales between host and symbiont may be beneficial for hosts if these differences enable them to acquire symbionts adapted to their settlement environments.

## 1. Introduction

The spatial scale of genetic variation differs widely between species, from globally panmictic dragonflies to tuataras that exhibit genetic structuring on the sub-kilometer scale (Moore et al., 2008; Troast et al., 2016). These contrasting patterns stem partially from differences in dispersal. Longer dispersal distances can lead to higher connectivity and more homogeneity between sites, blunting the effects of drift and inbreeding for smaller populations. Shorter dispersal may enable spatial heterogeneity and local adaptation as populations evolve to tolerate their immediate environment (Slatkin, 1987). However, patterns of genetic variation may also be driven by co-dispersal among closely associated species, such as mutualistic pairs or host-microbe holobionts. How dispersal shapes landscape genetic structure in such associations remains poorly understood (Hand et al., 2015).

Some theorists have suggested that the evolution of mutualisms should rely heavily on parent to child transmission of symbiotic partners, and therefore codispersal of the mutualist pair (Ewald, 1987). This is certainly true in some cases. A wide variety of terrestrial plants, for example, transmit their mycorrhizal partners directly from parent to offspring (Johnston-Monje et al., 2021; Laurent-Webb et al., 2024). However, the real world provides contradictory examples, including the many species of corals that acquire their algal obligate symbionts environmentally, a situation that some authors have described as a “paradox” (Hartmann et al., 2017). Environmental acquisition of mutualist symbionts occurs across taxa, and is especially common in the marine realm, including in many species of corals (Russell, 2019). Photosynthetic corals play critical roles in maintaining global biodiversity and human wellbeing, and depend on algal symbionts for their nutrition and survival (Fisher et al., 2015; Mellin et al., 2022). While corals and symbionts form obligate mutualist associations, they can disperse separately, making them an ideal study system to understand the extent to which mutualists show similar patterns of dispersal.

Most reef-building corals depend on photosynthetic algae living within their tissues to produce up to 95% of the carbohydrates they need to survive (Muscatine & Cernichiari, 1969). These algal symbionts belong to multiple genera, and individual corals can acquire symbionts in one of two ways. “Brooding” species of corals transmit their symbionts vertically from parent to offspring, producing larger larvae that carry their own symbionts. By contrast, “broadcast” spawning corals release gametes directly into the water column, where they form smaller larvae that acquire symbionts from the environment post-settlement (Harii et al., 2009). The second life history strategy, employed by corals of the genus *Acropora*, leads to the potential for corals and their algal symbionts to exhibit different patterns of genetic variation across space (Boutet & Schierwater, 2021). However, the extent to which corals and their symbionts show similar or disparate patterns of dispersal remains unclear.

Fortunately, population genetic methods allow direct observation of patterns of spatial genetic variation, as well as indirect inference about dispersal through isolation by distance methods (Pinsky et al., 2017). For example, analyses of *Stylophora pistillata* and *Pocillopora verrucosa* on the Great Barrier Reef found far higher cross-reef connectivity and genetic homogeneity in the latter species, with estimated mean dispersal distances of 2100 to 5200 meters in *P. verrucosa* but just 23 to 102 meters in *S. pistillata* (Meziere et al., 2025). Algal symbionts of corals may or may not show the same level of connectivity as their hosts. Some evidence shows algal symbionts exhibiting genetic differentiation at finer spatial scales than their coral hosts (Davies et al., 2020). This could imply consistent differences in both connectivity and local adaptation between corals and their symbionts.

Our limited understanding of how landscape genetic patterns differ between corals and their algal symbionts is especially critical for corals of the genus *Acropora*. This keystone genus of branching corals associates with a variety of algal symbiont strains acquired environmentally after their larval dispersal phase (Cumbo et al., 2013). All species of *Acropora* are broadcast spawners, and their asymbiotic larvae drift in the water column for days to weeks before settling (Nishikawa et al., 2003). *Acropora* in the Indo-Pacific, including the Philippines, mostly associate with symbionts from genus *Cladocopium*, though they can also associate with symbionts of other genera (Davies et al., 2020; Matias et al., 2023; Matsuda et al., 2022; Torres et al., 2021). *Acropora* are sensitive to heat waves and face a growing threat from anthropogenic climate change (Pratchett et al., 2013). In addition, some *Acropora* populations exhibit local genetic adaptation to higher temperatures at both the regional and intra-reef scales, as do their algal symbionts (Howells et al., 2012; McClanahan et al., 2020; Thomas et al., 2018). Consequently, understanding *Acropora* landscape genetics matters for applied conservation questions concerning connectivity, reserve design, and adaptation to anthropogenic change (Arafeh-Dalmau et al., 2023).

Here, we examine landscape genomic patterns of *Acropora cf. tenuis* and its algal symbionts in the central Philippines to test whether host and symbiont exhibit congruent or disparate dispersal patterns. The samples for this study are from the Philippines, which is situated within the Coral Triangle, known as the global epicenter of marine biodiversity (J. (Charlie) E. N. Veron et al., 2011). Our spatially dense sampling took place along two different two hundred kilometer stretches of reefscape on Cebu and Leyte islands, which facilitated assessment of isolation by distance patterns. Prior study of multiple species of reef-dwelling anemonefish across this same seascape found clear isolation by distance patterns and estimated dispersal kernel spreads on the order of 10 kilometers (Catalano et al., 2021; Fitz et al., 2023; Pinsky et al., 2010). Here, we test for similar patterns in *Acropora cf. tenuis* and its algal symbionts. Specifically, we examined whether (1) specific algal symbiont genotypes associated with specific *A. cf. tenuis* host genotypes and (2) host and symbionts showed similar or contrasting degree of genetic isolation by distance. Ultimately, we aimed to unpack the interacting relationships between genetic variation in corals, genetic variation in their algal symbionts, and physical location in space.

## 2. Materials and methods

### 2.1 Field sampling, field identification, and genomic library preparation

Field sampling was conducted from January to April 2016 across 47 sites in the Visayas region of the central Philippines, including areas surrounding the Camotes Sea and Cebu Strait. Sites were distributed across Cebu Province, including Cebu Island (n = 27 sites) and the Camotes Islands (n = 5), as well as Leyte Province (n = 15). Sites were selected to be approximately 10 km apart to capture ecologically relevant dispersal distances and enable testing for isolation by distance patterns in the study region (Fig. 1).

**Fig. 1.**
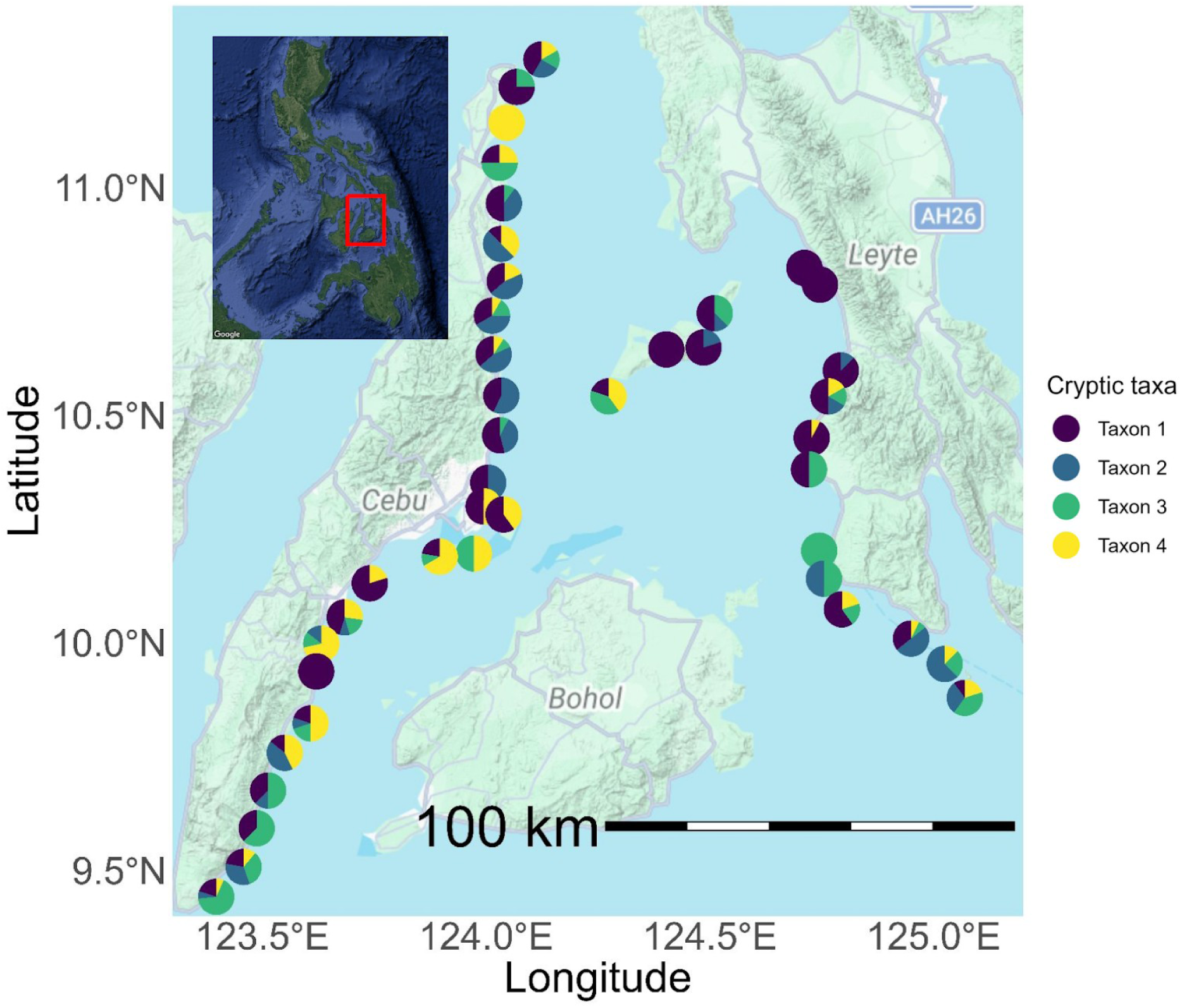
Map of sampling sites in the central Philippines. The red rectangle in the inset map represents that area of the primary map. Each pie chart represents a separate sampling site. Pie charts show the percentage of *Acropora* samples collected at each site assigned to each cryptic taxon. Some sites with only one taxon present had only one or two total samples (see Supplemental Table 1 for sample sizes). While taxa occurrence is weakly structured by latitude, all genetic groupings occurred at multiple sites on Cebu, Leyte, and the Camotes Islands.

At each site, we characterized benthic cover using line point intercept transects. Three 15 m transects were deployed haphazardly parallel to shore, and the bottom type was recorded at 20 cm intervals along each transect. Following benthic surveys, we collected tissue samples from *Acropora cf. tenuis* colonies at each site. To locate colonies, we conducted a separate belt transect survey at each site by swimming parallel to shore within a 2 m wide belt for approximately one hour or until 20 samples were collected. During these surveys, we measured the maximum length, width, and depth of every *A. cf. tenuis* colony encountered and recorded the time of collection. We collected small tissue samples (approximately 2 cm × 2 cm) from 20 colonies per site, ensuring that sampled colonies were spaced at least 10 m apart to avoid resampling the same genet. A towed GPS unit was used to record the coordinates of each sampled colony. If fewer than 20 samples were obtained during the initial survey, an additional dive was conducted to complete the target sample size (depth information was not collected during these additional dives).

*Acropora cf. tenuis* specimens were identified in the field based on morphological characteristics described in Veron’s “Corals of the World” (J. E. N. Veron, 2000). Key characteristics used for identification included: 1) colony growth form characterized by corymbose clumps, 2) long and tubular axial corallites, and 3) neatly arranged and slightly flaring radial corallites. Field identification focused on the combination of branching pattern, corallite structure, and overall colony architecture rather than relying on any single morphological trait. Detailed close-up photographs were taken of each collected specimen and are available at https://doi.org/10.5061/dryad.sf7m0cgnr.

In total, 258 samples were collected during initial surveys and 394 samples on additional dives. Tissue samples were preserved in 95% ethanol for genetic analysis. All samples were collected under Gratuitous Permit 0129-17 and transported to the United States under CITES export permit 4733. The Texas A&M Corpus Christi Genomics Core Laboratory performed all DNA extraction using silica column membrane kits, as well as double digest restriction-site associated DNA sequencing (ddRadSeq) library preparation (Peterson et al., 2012), and sequencing. Library preparation for ddRadSeq used the enzymes SphI and EcoRI and targeted fragment sizes of 500-600 basepairs. All sample libraries were pooled and then sequenced across two Illumina HiSeq4000 2 x 150 bp lanes to mitigate the possibility of variation introduced by lane effects.

### 2.2 Sequence processing, alignment, and filtering

We did initial quality control and 3’ read trimming with FastP version 0.23.4 (Chen et al., 2018), visualized the results with MultiQC (Ewels et al., 2016), de-duplicated and compressed the read files with the Clumpify function from BBtools version 39.06, re-ran FastP for 5’ read trimming, and re-paired unpaired reads with BBtools (Bushnell, 2014). We parallelized analysis with GNU Parallel version 20200122 (Tange, 2018). Full scripts with parameters are available on Github at https://github.com/pinskylab/Atenuis_Philippines (to be replaced with a Zenodo DOI upon manuscript acceptance). The pre-processing scripts were developed from Bird et al. (2024). We then mapped the trimmed reads to genomes from *Acropora tenuis* (Shinzato et al., 2021), *Cladocopium goreaui* (Martin-Cuadrado et al., 2024), *Durusdinium trenchii* (Dougan et al., 2024), and *Symbiodinium kawagutii* (Martin-Cuadrado et al., 2024), all downloaded from NCBI, using BWA version 0.7.17 (Li & Durbin, 2009). The *‘Acropora tenuis*’ genome assembly available on NCBI comes from Japan and is therefore unlikely to represent true *A. tenuis* (Bridge et al., 2024). Given the broad taxonomic uncertainty and extensive presence of cryptic species within *Acropora cf. tenuis*, this genome likely represents a congener to our samples, rather than a conspecific. Across individuals, a median of 41,236.5 reads, or 74% of total reads, mapped to the *Acropora tenuis* genome from NCBI. A median of 14,424.5 reads, or 15% of the total, successfully mapped to the *Cladocopium goreaui* genome, 2771 reads (2.4% of total) mapped to the *Symbiodinium kawagutii* genome, and 1447 reads (1.5% of total) mapped to the *Durusdinium trenchii* genome.

After sorting and indexing the resulting bamfiles with Samtools version 1.20 (Li et al., 2009), we called SNPs using FreeBayes version 1.3.10 (Garrison & Marth, 2012), resulting in 2,491,826 *Acropora* SNPs across 652 individuals and 596,153 *Cladocopium* SNPs across 644 individuals. Nearly all individuals showed substantial missing data, with 184 (28%) of individuals showing more than 99.9% missing data after selecting only loci with a minimum quality score of >30 and a minimum depth of >3. More than half of individuals (359, 55%) were missing more than 90% of possible SNPs. We iteratively filtered SNPs and individuals using functions from vcftools and vcflib (Danecek et al., 2011; Garrison et al., 2022), in order to preserve the maximum number of each, as described in O’Leary et al. (2018).

Ultimately, we retained all SNPs present in at least 85% of individuals and all individuals with at least 50% non-missing data. After quality filtering, we were left with 2,269 SNPs and 301 individuals in the *Acropora* assemblies, and 2,059 SNPs and 310 individuals in the *Cladopium* assemblies. We pruned SNPs for physical linkage using Plink2 by examining SNPs across a 50 bp window size and pruning those with r^2^>0.5 (Chang et al., 2015). This process left us with 1,838 SNPs in *Acropora* and 873 SNPs in *Cladocopium*.

We calculated distance to shore for each sample using the global shorelines dataset from OpenStreetMap, downloaded on March 28, 2025 (OpenStreetMap contributors, 2025).

### 2.3 Identification of genetic clusters within *Acropora*

*A. tenuis* was traditionally believed to be a common coral species across shallow reefs in the western Pacific ocean and Red Sea. However, recent research has revealed it to comprise at least eleven morphologically similar but evolutionary distinct species (Bridge et al., 2024; J. E. N. Veron, 2000). Multiple species of *A. tenuis* lookalikes have been identified co-occurring on the same reefs in Australia, Okinawa, and the South Pacific (Bridge et al., 2024; Matias et al., 2023; Zayasu et al., 2021). While some of these *A. cf. tenuis* populations have been provisionally reassigned to new nominal species, those in Southeast Asia (including the Philippines) have so far only been defined as *Acropora sp(p).* The authors of a recent taxonomic revision note that this should be considered “an unresolved clade requiring further sampling and analysis” (Bridge et al., 2024). Here we refer to our samples as *Acropora cf. tenuis* to reflect both the species’ traditional classification and current taxonomic uncertainty.

We determined the number of genetic clusters within the fully filtered *Acropora* SNP data using a Discriminant Analysis of Principal Components (DAPC) including the top 500 principal components, to capture the maximum amount of variation, as recommended by the package authors (Jombart & Collins, 2015). We determined the optimal solution by comparing Bayesian Information Criteria (BIC) for all solutions between one and sixteen clusters. We calculated principal components eigenvectors and eigenvalues using Plink2 version 2.00a5.10 in order to visualize these clusters in PCA space and quantify the percent of genetic variation accounted for by each principal component. We also examined the top three principal components for linear correlation with latitude, longitude, distance to shore, and depth.

We quantified genetic distance between clusters by calculating FSTs between groups using the Hierfstat R package version 1.1.10 (Goudet, 2005). We then reshuffled all individuals between each pair of taxa and recalculated the FST 1000 times in order to simulate a null distribution of genetic distances. To assign statistical significance to the pairwise genetic distance between clusters, we compared each observed between-group FST to its simulated null distribution and calculated the percentile of the null distribution greater than the observed FST.

We also evaluated ancestry for each individual sample using ADMIXTURE version 1.3.0 and all possible numbers of ancestry groups between one and sixteen using a cross validation procedure (Alexander et al., 2009). This approach found ancestry groups that corresponded closely to the genetic groups found by the DAPC. Given these consistent differences and the substantial genetic differentiation between these groups despite their broadly overlapping ranges (see Results), we referred to these groupings as “taxa,” which we numbered arbitrarily one through four with the most individuals assigned to cluster one and the fewest assigned to cluster four. We used the DAPC to assign each *Acropora* individual to a genetic cluster, and re-filtered SNP data separately within each cluster (Section 2.2). We used slightly different iterations of SNP and individual removal, but with the same final cutoffs as described above. After pruning, we ended with 116 individuals and 2308 SNPs in Taxon 1, 69 individuals and 1689 SNPs in Taxon 2, 57 individuals and 1221 SNPs in Taxon 3, and 53 individuals with 1213 SNPs in Taxon 4. We identified potential clones using the KING coefficient calculated in Plink2 using a cutoff of 0.354 (McMaster et al., 2025). We calculated heterozygosity and inbreeding coefficients (F_IS_) for each individual sample and averaged across taxa using vcftools.

### 2.4 Genetic variation with environment and geography in *Acropora*

Non-parametric Kruskal-Wallis tests in R were used to test for differences in mean latitude, depth, and distance to shore between genetic clusters. We calculated isolation by distance (IBD) patterns for each genetic group individually in accordance with Sheets *et al*. (2018). We assessed isolation by distance in both one dimension (along the coast of Cebu) and two dimensions (all sites together). We did not analyze one dimensional patterns along the coast of Leyte because of insufficient coral samples along that coastline. We calculated pairwise linearized FST and the pairwise geographic distance between each pair of sampling sites. Only sites with at least two individuals were considered for FST calculations. For the two dimensional IBD assessment, we took the natural logarithm of these distances, following the equation for isolation by distance in two dimensions (Rousset, 1997). Mantel tests were conducted comparing the genetic and geographic distance matrices using the Vegan package version 2.6.8 (Dixon, 2003). In order to compare strength of IBD relationships between taxa, we calculated confidence intervals around regression slopes by using the standard error of the regression slope as the standard deviation for 1000 normally distributed slopes centered on the estimated regression coefficient. We then quantified the percentage of these resampled regression slopes that were greater than zero, indicating a positive relationship between geographic and genetic distance. Finally, we identified putative parent/offspring pairs from *Acropora* SNP data using the GetMaybeRel() function of the Sequoia package version 3.0.3 in R with no age priors, assuming all individuals as hermaphrodites (Huisman, 2017).

### 2.5 Taxonomic variation within algal symbionts

To identify the taxonomic composition of the symbionts in our samples, we compared the percent of reads mapped to genomes from three common genera of algal symbionts (*Cladocopium goreaui, Durusdinium trenchii,* and *Symbiodinium kawagutii*). To identify environmental correlates of symbiont composition, we used Kruskal-Wallis tests to compare the relative percentages of reads mapped from each individual to each genus to depth, distance to shore, latitude, island, and host coral genetic grouping. In nearly all individuals, by far the most reads mapped to the *Cladocopium goreaui* genome, leading us to proceed with further analysis for only these assemblies (see Supplemental Fig. 2 for percent of reads mapping to each symbiont genome).

Similarly to *Acropora*, we used a DAPC and paired PCA to evaluate the number of genetic clusters and to visualize genetic groupings in PCA space. We found only one genetic cluster, and consequently did not subdivide our *Cladocopium* population for subsequent analyses.

We also ran an ADMIXTURE analysis for the *Cladocopium* genetypes, again evaluating all possible numbers of ancestry groups between one and sixteen using cross-validation. We assigned *Cladocopium* genomes to groups based on the taxon (1- 4) of their *Acropora* host, calculated FSTs between those taxa, and calculated associated p-values using the same resampling procedure for simulated null distributions described in Section 2.4. After SNP filtering, more individual samples were found with >50% data present for the *Cladocopium* assemblies than for the *Acropora* assemblies, meaning that some (n=10) *Cladocopium* samples could not be assigned to *Acropora* taxa.

### 2.5 Isolation by distance and genetic variation with environment in *Cladocopium*

We used linear regression to compare the top three principal components of the *Cladocopium* genomes to latitude, longitude, distance to shore, and depth. We assessed isolation by distance in both one and two dimensions using a Mantel test from the Vegan package in R following the same methodology as described in Section 2.4. Confidence intervals around slopes were calculated similarly to the *Acropora* samples. To account for differences in sample size and geographic distribution of samples between the *Acropora* clusters and the full set of *Cladocopium* samples, we also evaluated isolation by distance patterns for *Cladocopium* within each genetic grouping of *Acropora* and reported distributions for those slopes as well. Finally, we used two partial Mantel tests to evaluate the influence of geographic distance on genetic distance between *Cladocopium* populations while controlling for genetic distance between *Acropora* populations, as well as the influence of genetic distance between *Acropora* populations on the genetic distance between *Cladocopium* populations while controlling for geographic distance. We did not test for kin pairs within the *Cladocopium* samples, as they are haploid (Santos & Coffroth 2003). All analyses using R packages were performed using R version 4.4.1 and all Python analyses were performed in Python 3.9.25.

## Results

### 3.1 Genetic groups in *Acropora cf. tenuis*

Our analysis revealed four distinct genetic clusters within the *Acropora cf. tenuis* genetic data. The four cluster solution resulted in the lowest BIC for the DAPC, closely followed by the five cluster solution (Supplemental Table 2). These clusters appear distinct in PCA space, with more than 50% of the genetic variation loading on the first two principal components. However, some intermediate individuals may represent hybrids (Fig. 2A). Likewise, the ADMIXTURE analysis with four ancestry groups resulted in the lowest cross validation error, with the five ancestry groups in second place (Supplemental Table 2). Most individuals showed a clearly dominant ancestry group, with 99% of individuals having more than half of their ancestry coming from a single group and 90% of individuals having at least 7/8th of their ancestry from a single group (Fig. 2B).

**Fig. 2.**
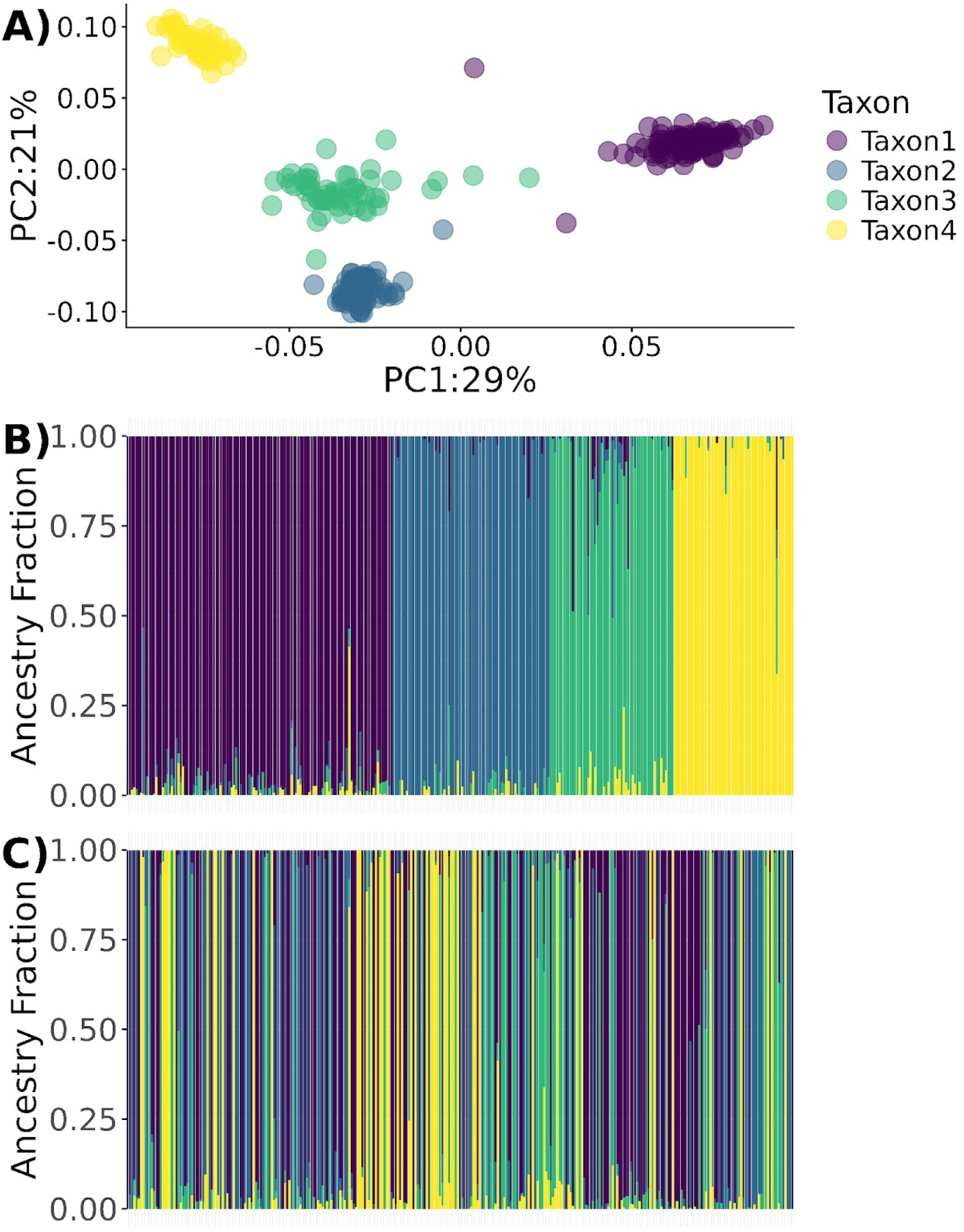
*Acropora cf. tenuis* genetic samples belong to four distinct clusters. **A.** A Discriminant Analysis of Principal Components resolved four clusters, which we color and then plot on PCA axes one and two. PC1 accounted for 29% of the variation, and PC2 accounted for 21%. **B.** An ADMIXTURE analysis found four ancestry groups, and most individual samples had the majority of their ancestry derived from a single group. Individuals in this plot are grouped horizontally according to which ADMIXTURE group supplies the largest fraction of their ancestry. **C.** Sorting the ADMIXTURE plot by latitude shows intermixing of ancestry groups along a latitudinal gradient.

The ancestry groups found with the ADMIXTURE analysis corresponded well but imperfectly to the genetic clusters resolved by DAPC. The vast majority (298/301, 99%) of individuals assigned to a cluster by DAPC had greater than 1/2 ancestry fraction coming from that same cluster, and 90% (271/301) had at least 7/8th of their ancestry from that cluster.

Pairwise FST comparisons between these genetic clusters ranged from 0.10 to 0.16 (Table 1). For each pairwise comparison, the actual FST was higher than 100% of the simulated resamples with shuffled individuals.

**Table 1.** FSTs between cryptic taxa and associated p-values. All FSTs were higher than 100% of those on a null distribution calculated by reshuffling individuals between taxa.

|  | Taxon 2 | Taxon 3 | Taxon 4 |
| --- | --- | --- | --- |
| Taxon 1 | $F_{ST}=0.12$<br>$p=0.00$ | $F_{ST}=0.11$<br>$p=0.00$ | $F_{ST}=0.16$<br>$p=0.00$ |
| Taxon 2 | | $F_{ST}=0.10$<br>$p=0.00$ | $F_{ST}=0.15$<br>$p=0.00$ |
| Taxon 3 | | | $F_{ST}=0.12$<br>$p=0.00$ |

These four genetic clusters were not separated in space. Each cluster occurred in at least 60% (26/43) of sampling sites and on both islands (Fig. 1). Taxa did appear somewhat structured by latitude (Kruskal-Wallis test, p-value=0.00026, n=301), with Taxon 2 having the highest median latitude (10.59°N) and Taxon 3 having the lowest (10.00 °N). There was a marginally significant association between genetic grouping and average depth (Kruskal-Wallis test, p-value=0.07, n=97), with Taxon 1 having the greatest depth (13 ft) and Taxon 4 having the lowest median depth (9.5 ft). There was no association with distance to shore (Kruskal-Wallis test, p-value=0.22, n=301).

We found no association between the number of genetic reads mapped per sample and taxon (Kruskall-Wallis test p=0.22, n=301). Taxa did show significant differences in inbreeding coefficient (F) (Kruskall-Wallis test p=3.1 x 10 ^−6^, n=301). The median inbreeding coefficient for Taxon 4 was negative (F=-0.023) indicating observed homozygosity slightly lower than expected. Inbreeding coefficients were positive but low (F=0.018 and F=0.0011) for Taxa 1 and 2, and higher in Taxon 3 (F=0.076). See Supplemental Fig. 3 for boxplot of inbreeding coefficients.

### 3.2 Isolation by distance and dispersal in *Acropora cf. tenuis*

For two out of four taxa, there was on average a positive correlation between two-dimensional geographic distance between populations in log meters and genetic distance in pairwise linearized FST. However, the correlations for Taxa 1 and 2 were negative, and all confidence intervals from resampling slopes overlapped with zero (Fig. 3). Moreover, none of the relationships between genetic distance and geographic distance were statistically significant (p<0.05) when using a Mantel test (Supplemental Tables 6 and 7). The strongest relationship, found in Taxon 4, was largely driven by three closely related individuals collected from sites at the north and south ends of Cebu, and excluding these individuals substantially reduced the percentage of positive IDB slopes in 1D for this taxon (Table 3). These individuals collectively account for the high FST values seen in some pairwise comparisons (Fig. 3G, H). Overall, a lack of strong IBD patterning suggests a high degree of genetic mixing within each cryptic *Acropora* taxon.

**Fig. 3.**
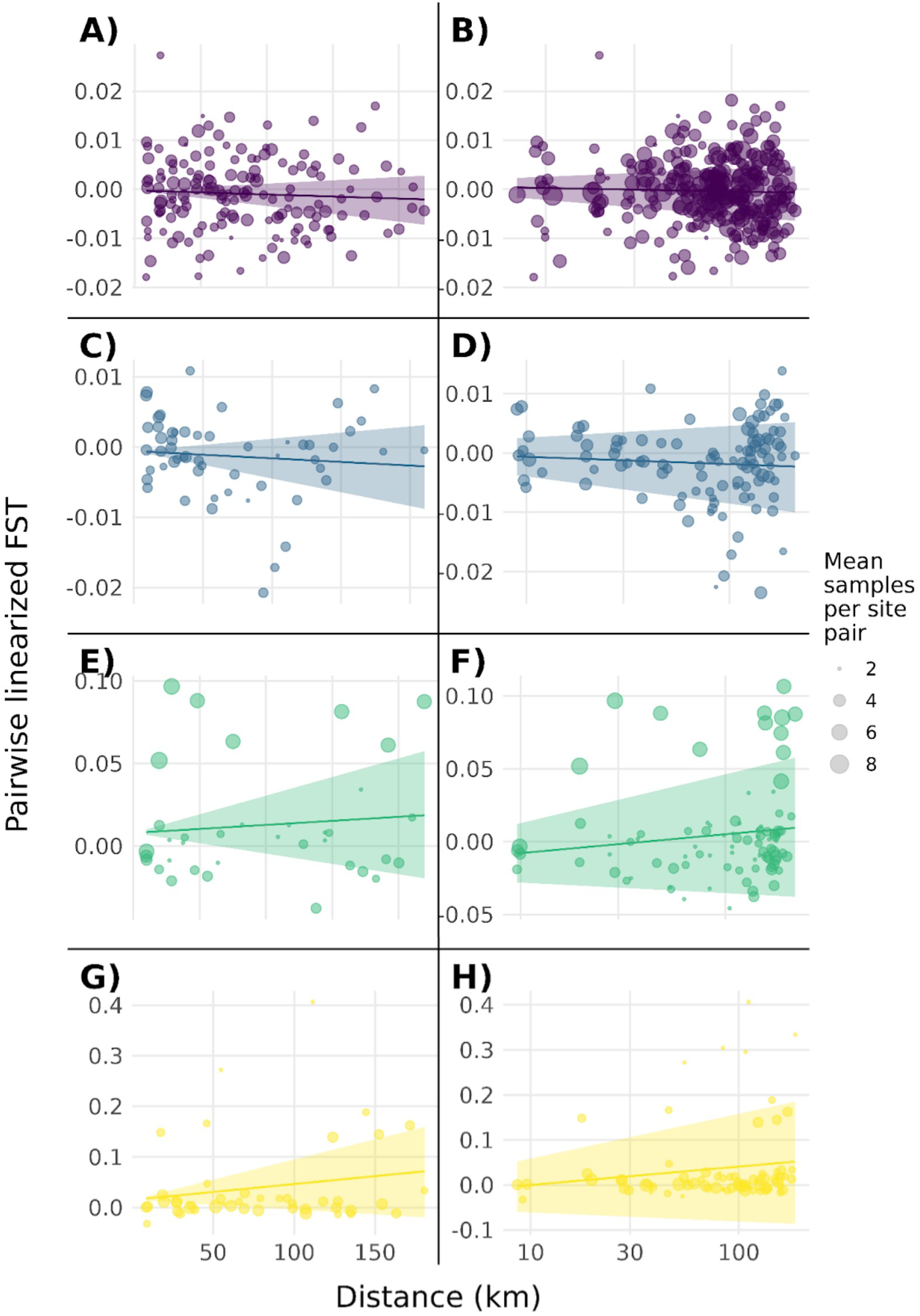
No *Acropora cf tenuis* taxon showed significant isolation by distance (IBD) patterns in either one or two dimensions. Each row shows one cryptic *Acropora* taxon (Taxon **1** - Taxon 4), plots on the left show one dimensional IBD comparisons using data from along the coast of Cebu only, and plots on the right show two dimensional IBD comparisons using data from all sites. Points correspond to pairwise comparisons of linearized FST and geographic distance (left panels) or log geographic distance (right panels) between sites. Shaded areas show the 95th percentile confidence intervals for the IBD slopes.

Pedigree analysis within the cryptic *Acropora* taxa found five putative parent/offspring pairs, or seven if both putatively clonal individuals at site CEB_01 were included (Supplemental Table 8). All parent/offspring pairs were genotyped at a minimum of 1000 of the same SNPs and showed a minimum log likelihood ratio of 4, indicating that the paired individuals were 10 ^4^ times more likely to represent parent/offspring pairs than they were to represent unrelated individuals. Three of these pairs were defined as clones in the original clone scan with KING, and are marked in Fig. 4 with a letter C. Nearly all of the pairs occurred at widely spaced sites (Fig. 4). These results suggest that long (>50km) dispersal distances are possible across a single generation.

**Fig. 4.**
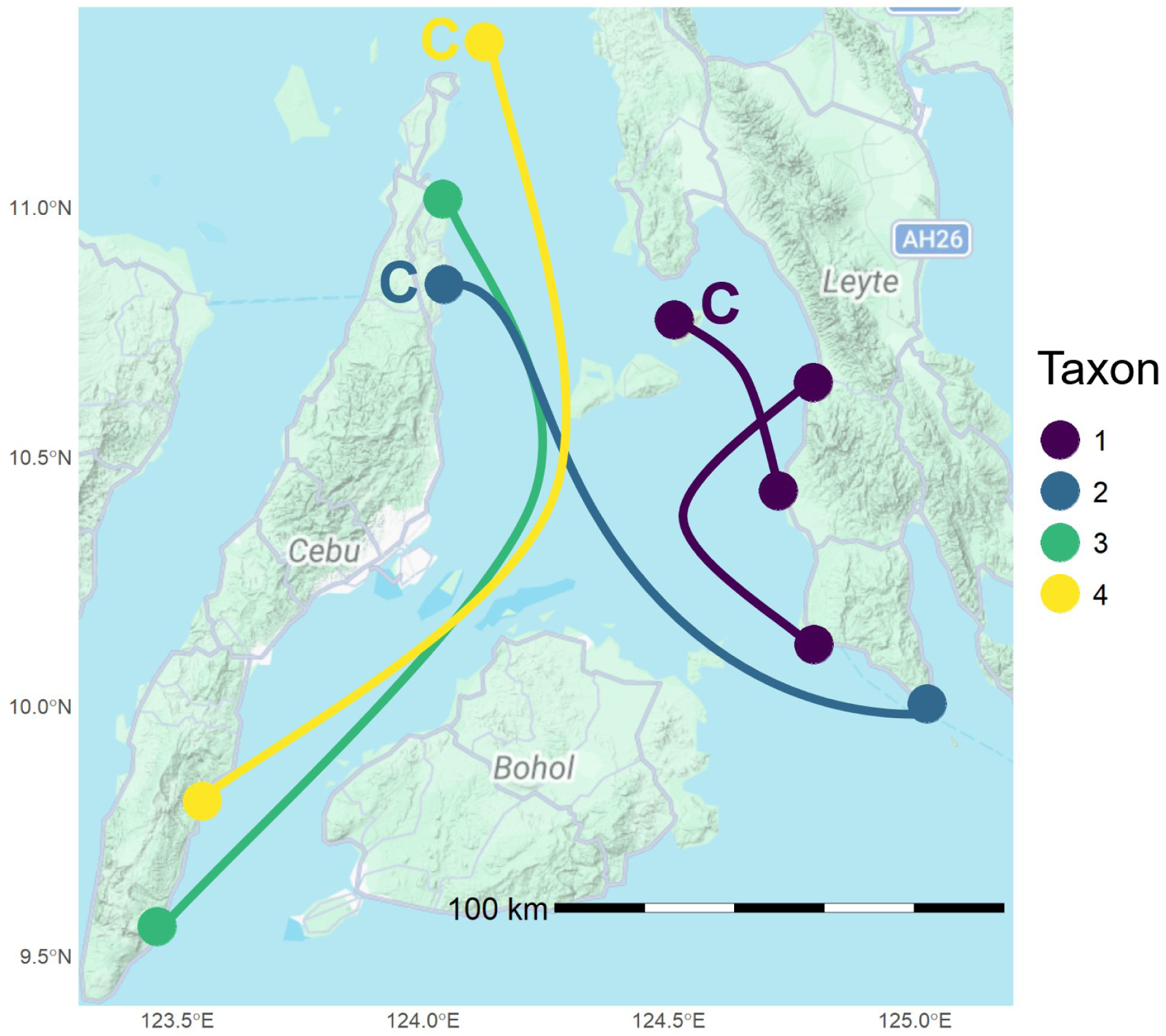
Putative parent/offspring pairs found by Sequoia exist at widely spaced sites. Dots show sites with samples included in parent/offspring pairs,and curves show connections between sites. Closely related individuals that may constitute either first degree relatives or clonal genotypes are marked with a letter C.

### 3.3 Identification of algal symbionts

We identified the vast majority of algal symbionts as the genus *Cladocopium* across all four *Acropora* taxa and 43 sites. More than 98% (644/652) of individual samples had a higher number of reads successfully map to the *C. goreaui* genome than either the *D. trenchii* or *S. kawagutii* genomes. Three individuals, two of which belonged to Taxon 1 and one of which belonged to Taxon 4, had the highest number of reads map to *D. trenchii.* One *Acropora* individual, which belonged to Taxon 1, had the highest number of reads map to *S. kawagutii* (Supplemental Tables 3 and 4). There was no significant relationship between percent of reads mapped to a given genus of symbiont, and host *Acropora* taxon (n=298, all p>0.5). Likewise, there was no significant association between relative number of reads mapping to different symbiont genera and either depth (Kruskal-Wallis test, p=0.71, n= 97) or distance to shore (Kruskal-Wallis test, p=0.13, n=295). Finally, linear regression models for each symbiont genus found no association between percentage of symbiont reads mapping to that genus and sample depth (n=97, all p>0.3). Since *Cladocopium* was by far the dominant symbiont genus for this dataset, we proceeded analyzing only symbiont DNA mapped to the *C. goreaui* genome. Given the low probability of our samples belonging to *C. goreaui* specifically, we refer to reads mapped to the *C. goreaui* genome by the more general term *Cladocopium*.

### 3.4 Genetic variation within *Cladocopium* and its correspondence to coral host genotype

*Cladocopium* did not show the same genetic groupings as their *Acropora* hosts. Principal components and discriminant analysis of principal components did not reveal distinct genetic clusters within the *Cladocopium* SNP profiles (Fig. 5A). The BIC for the DAPC was minimized by finding a single cluster, and BIC increased monotonically with the number of clusters up to at least 16 clusters (Supplemental Table 5). ADMIXTURE resolved three ancestry groups within the *Cladocopium*, but these ancestry groups did not obviously correspond to those found in their *Acropora* hosts (Fig. 5B). However, there was a noticeable structuring of ancestry with latitude (Fig. 5C).

**Fig. 5.**
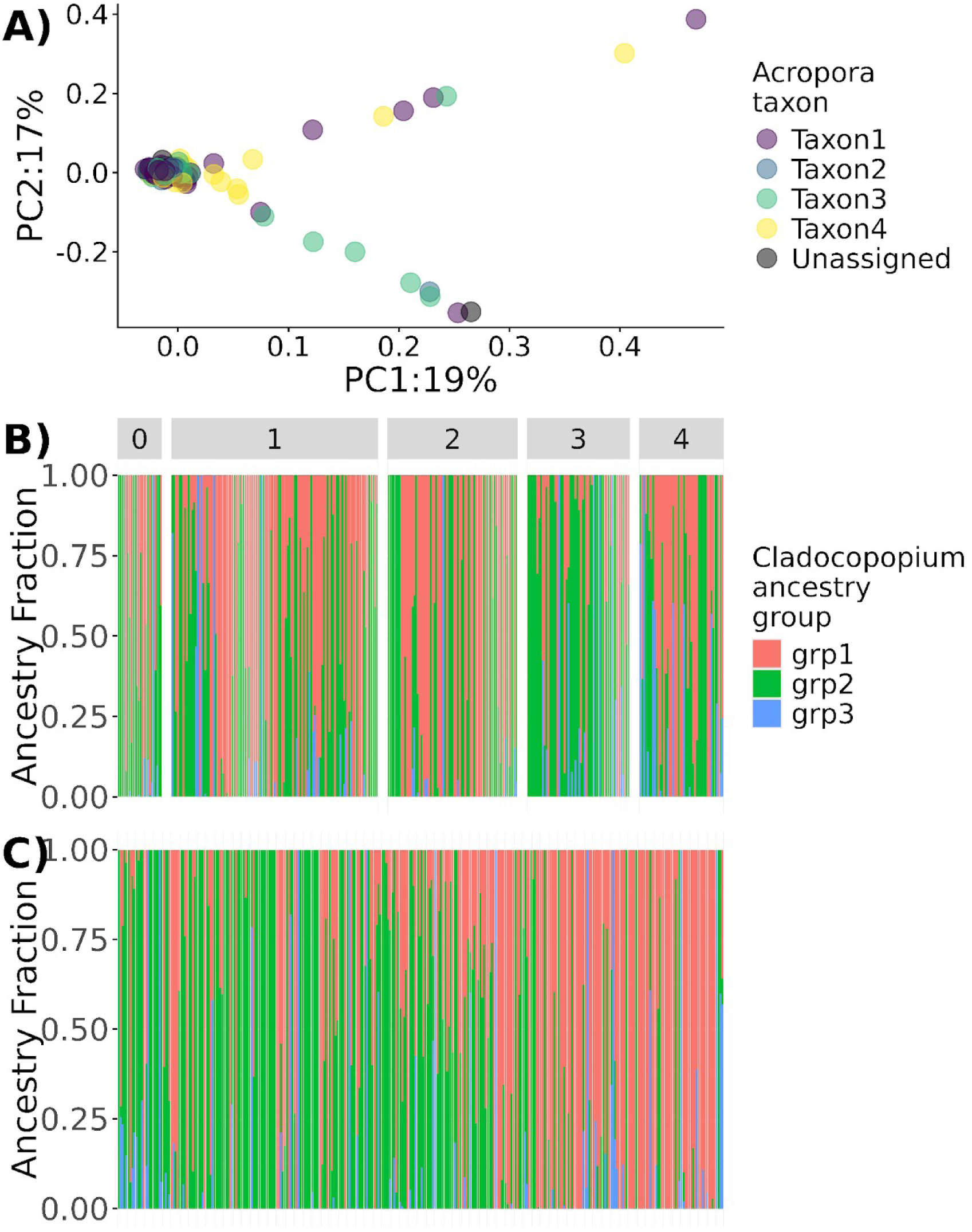
*Cladocopium* genetic samples all belong to the same genetic cluster. **A.** A Discriminant Analysis of Principal Component analysis resolved only one cluster. We color *Cladocopium* samples by host *Acropora* cryptic taxon and then plot on PCA axes one and two. PC1 accounted for 19% of the variation, and PC2 accounted for 17%. **B.** An ADMIXTURE analysis found three ancestry groups, and many individuals had mixed ancestry from multiple groups. Individuals are organized by taxon of *Acropora* host, as shown by the numbered gray bars. Individuals labeled as ‘0’ belong to hosts not assigned to a taxon due to insufficient reads of *Acropora* DNA. **C.** Sorting the ADMIXTURE plot by latitude revealed differences in *Cladocopium* ancestry with latitude.

Based on this PCA, *Cladocopium* populations do show evidence for genetic structuring in space across both regional and microhabitat scales. The second principal component of the *Cladocopium* genomes had a significant linear relationship to latitude (linear regression, p=0.00027, n=310). Similarly, there is a significant relationship between the first principal component of the *Cladocopium* genomes and host coral depth (p=0.000013, n=107).

*Cladocopium* populations associated with different *Acropora* cryptic taxa showed weak differentiation with extremely low FSTs. However, reshuffling individuals between taxa and recalculating FSTs of the associated *Cladocopium* populations reveals that while weak, some of these FSTs were significantly stronger than would be expected from random association (Table 2).

**Table 2.** FSTs between *Cladocopium* populations associated with different cryptic taxa of *Acropora* are extremely low, yet on average somewhat higher than would be expected by chance. The difference between those *Cladocopium* in Taxon 1 and those in Taxon 3 is particularly significant.

|  | Taxon 2 | Taxon 3 | Taxon 4 |
| --- | --- | --- | --- |
| Taxon 1 | $F_{ST}=0.00069$<br>$p=0.236$ | $F_{ST}=0.0047$<br>$p=0.000$ | $F_{ST}=0.00069$<br>$p=0.291$ |
| Taxon 2 | | $F_{ST}=0.0034$<br>$p=0.026$ | $F_{ST}=0.0040$<br>$p=0.013$ |
| Taxon 3 | | | $F_{ST}=0.0019$<br>$p=0.176$ |

### 3.5 Isolation by distance in *Cladocopium*

*Cladocopium* showed more consistent patterns of genetic isolation by distance than did their coral hosts. Mantel tests found significant positive associations between pairwise linearized FST and geographic distance in one dimension along the coast of Cebu (p=0.002, n=23 sites) and between pairwise linearized FST and log geographic distance in two dimensions across the entire study area (p=0.001, n=36 sites). A higher percentage of the resampled slopes (99.7%) were positive than they were for any of the *Acropora* taxa, and the 95% confidence interval did not overlap with zero (Fig. 6).

**Fig. 6.**
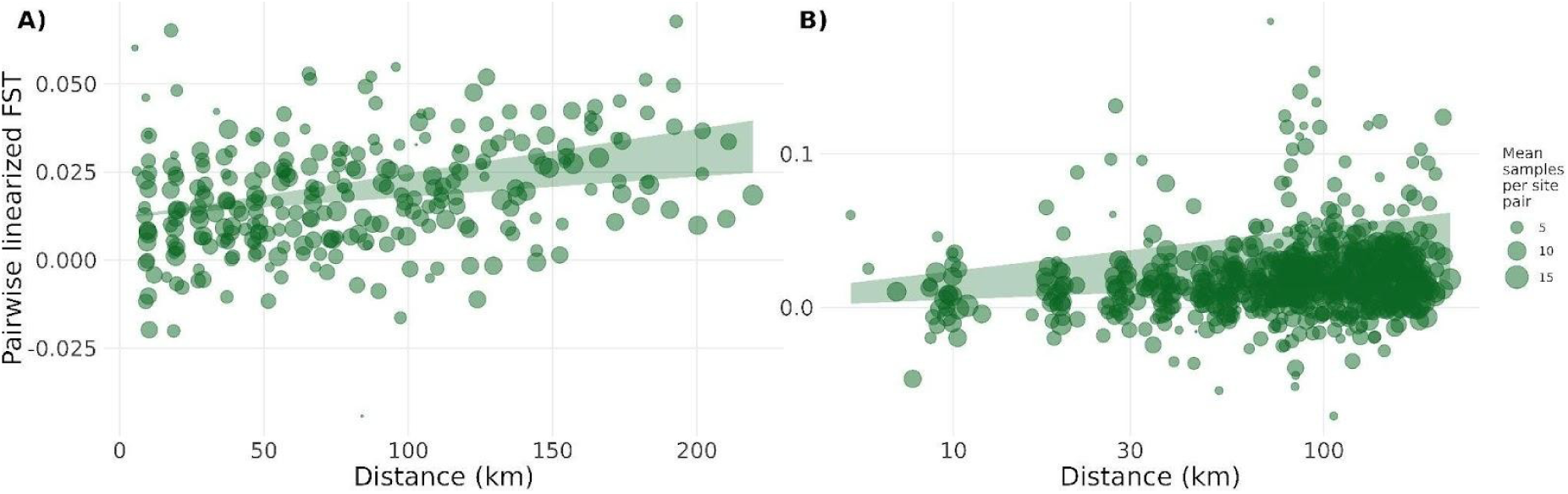
Mantel tests reveal consistent positive relationships between genetic distance in pairwise linearized FST and geographic distance in both (A) one and (B) two dimensions for *Cladocopium*.

When we analyzed IBD slopes for *Cladocopium* within each cryptic taxon separately to control for differences in sample size and geographic distribution of samples between *Cladocopium* and *Acropora* taxa, a higher percentage of IBD slopes were positive for the *Cladocopium* samples than for the *Acropora* samples in four out of four taxa for the one dimensional analysis, and in two out of four taxa for the two dimensional analysis (Table 3). In the two dimensional analysis, a higher percentage of IBD slopes were positive for three out of four *Cladocopium* samples than for the *Acropora* samples after excluding one potential clone. For Taxon 3, both *Cladocopium* and *Acropora* resampled IBD slopes were positive for a large fraction of the draws, with a slightly higher percentage in *Cladocopium* for one dimensional analyses, and slightly higher for *Acropora* for two dimensional analyses (Table 3).

**Table 3.**
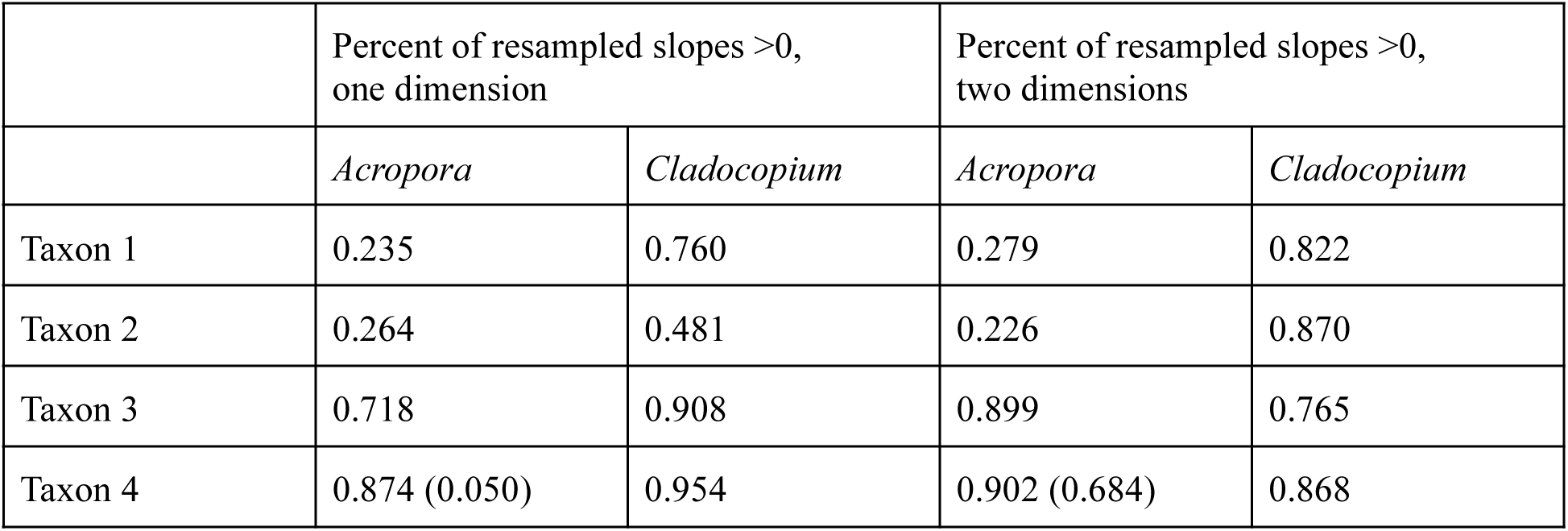
A higher proportion of resampled slopes are positive in *Cladocopium* than in their *Acropora* hosts when accounting for sample size and geographic distribution. Rows correspond to individual corals assigned to each cryptic taxon. The left two columns show one dimensional analysis along the coast of Cebu for *Acropora* genotypes and their associated *Cladocopium*, and the right two columns show two dimensional analysis across all sites. For the final row, the values in parentheses show the proportion of positive slopes when excluding one of a pair of putative clones that occurred at the same site.

|  | Percent of resampled slopes >0,<br>one dimension |  | Percent of resampled slopes >0,<br>two dimensions |  |
| --- | --- | --- | --- | --- |
|  | *Acropora* | *Cladocopium* | *Acropora* | *Cladocopium* |
| Taxon 1 | 0.235 | 0.760 | 0.279 | 0.822 |
| Taxon 2 | 0.264 | 0.481 | 0.226 | 0.870 |
| Taxon 3 | 0.718 | 0.908 | 0.899 | 0.765 |
| Taxon 4 | 0.874 (0.050) | 0.954 | 0.902 (0.684) | 0.868 |

Geographic distance had a stronger influence on *Cladocopium* genetic similarity than did host *Acropora* genotype. A partial Mantel test assessing the relationship of FSTs between *Cladocopium* populations and FSTs between *Acropora* populations conditioned on geographic distance was not significant (p=0.347, n=36 sites). Conversely, a partial Mantel test of between-population *Cladocopium* FSTs and geographic distance conditioned on *Acropora* between-population FSTs was significant (p=0.004, n=36 sites).

## Discussion

This study used dense genetic sampling across a seascape to infer patterns of dispersal and, more broadly, spatial genetic variation in *Acropora cf. tenuis* corals and their algal symbionts. Unexpectedly, we found four genetically distinct but sympatrically distributed groups of *Acropora* within our sample. All of the taxa exhibited genetic patterns across the study area consistent with long-distance dispersal, and we found putative parent/offspring pairs at widely spaced sites. However, all genetic clusters of *Acropora* hosted primarily a single type of algal symbiont belonging to the genus *Cladocopium*. *Cladocopium* genotypes were structured more strongly by geographic location rather than host genotype. When comparing dispersal across hosts and symbionts, the algal symbionts showed greater genetic structure in space than did their *Acropora* hosts.

### Sympatric cryptic diversity within *Acropora cf. tenuis*

Prior study of *Acropora cf. tenuis* suggests this species actually constitutes multiple morphologically similar species. In particular, the *Acropora cf. tenuis* group in Southeast Asia is poorly defined in terms of both genetics and morphology, and may constitute multiple lineages (Bridge et al., 2024). Our results strongly support the existence of at least four sympatric genetic groups within *Acropora cf. tenuis* in the Philippines, along with limited hybridization between those groups. Multiple analysis methods converged on four groups in our samples and assigned individual corals to groups with a high level of consistency, despite their lack of differentiation in space. Combining these lines of evidence, we consider these groups to be cryptic taxa, possibly constituting distinct members of a species complex. In particular, our ADMIXTURE results suggest limited interbreeding between the four ancestry groups, though the existence of mixed ancestry in some individuals implies that intermittent hybridization between these genetic clusters can occur.

Species definitions in *Acropora* are frequently ambiguous, with extensive hybridization between species (Ladner & Palumbi, 2012; Wu et al., 2024). Closely related *Acropora* species may be best defined as “syngameons,” or groups of two or more related species that are genetically distinct, yet regularly interbreed (Ladner & Palumbi, 2012). Given the apparently incomplete genetic separation of genetic groups within *A. cf. tenuis* individuals in our dataset, these populations may fit that definition. While prior studies of *Acropora* species complexes have found microhabitat differences associated with different cryptic taxa (Ladner & Palumbi, 2012; Matias et al., 2023), we found only weak evidence of microhabitat differentiation between the cryptic taxa found here, defined in terms of depth and distance to shore. However, we had incomplete data on individual coral depths, which may have hindered our ability to observe such differentiation. Our results underscore the need for further study of *Acropora* taxonomy in the Philippines, which would allow us to uncover potentially differing distributions, ecological roles, and conservation needs of these previously conflated taxa.

### Long-distance dispersal and weak spatial genetic variation in *Acropora*

Broadcast spawning corals, including *Acropora*, often show long distance connectivity through dispersal of their pelagic larvae. For example, a study of connectivity among *Acropora austera* populations in Mozambique found genetic near-panmixia across a 384 km stretch of coastline (Duvane et al., 2025). A genetic study of *A. hyacinthus* and *A. digitifera* in the south Pacific found isolation by distance patterns in both, but only on the spatial scale of thousands of kilometers (Davies et al., 2020). Research on *Pocillopora verrucosa*, a coral belonging to a different genus but with a similar reproductive mode to *Acropora*, on the Great Barrier Reef estimated dispersal distances anywhere between 21 and 52 kilometers using isolation by distance relationships to estimate dispersal kernel spread (Meziere et al., 2025).

Patterns of spatial genetic variation in each cryptic *Acropora* taxon studied here are consistent with long larval dispersal distances. We found no significant isolation by distance slopes in one or two dimensions despite dense sampling across 200 km. Confidence intervals for all slopes overlapped with zero, and resampling slopes resulted in substantial percentages with non-positive slopes. For one taxon (Taxon 2), the best-fit slope was less than zero. Taxon 4 also showed an average negative isolation by distance slope in the one dimensional analysis after excluding clones. All of this suggests minimal or no isolation by distance within the *Acropora* samples at 200 km scales. While still not significant, Taxon 3 had marginally stronger isolation by distance patterns than the other taxa, which may indicate shorter average dispersal in that taxon. However, this could also be due to lower effective population size, which would be consistent with the higher inbreeding coefficient in this taxon.

Unlike the results presented here, multiple studies of reef fishes in this same area of the Philippines have found clear isolation by distance patterns, which were then used to estimate dispersal kernel spreads. A 2010 study of anemonefish *Amphiprion clarkii* across Cebu and Leyte found a dispersal kernel spread of 11 km, with a confidence interval of 4 - 27 km (Pinsky et al., 2010), with additional research showing that dispersal varied both seasonally and interannually (Catalano et al., 2024). A study in the sister taxon *Amphiprion biaculeatus* found a similar dispersal kernel spread of 8.9 km, with a confidence interval of 2.3 - 18.4 km (Fitz et al., 2023).

The lack of any consistent isolation by distance pattern in *Acropora* across exactly the same geography as that used in the *Amphiprion* studies suggests longer, and perhaps much longer, dispersal distances for *Acropora* corals than their co-occurring reef fishes. Rousset (1999) recommends a square sampling area for isolation by distance patterns of at least 10 times the dispersal distance of the study organism. In the case of this approximately 200 km by 100 km area, this guideline would suggest that clear isolation by distance patterns would have been apparent for dispersal distances of less than ten to twenty kilometers. The lack of such patterns suggests that *Acropora* dispersal may regularly exceed this threshold.

Kinship analysis added further evidence to this hypothesis, finding closely related individuals of *Acropora* separated by as much as 181 km. While these distances are mostly long, they are not implausible given the oceanography of the area and the life history of broadcast spawning corals. Some of the distances between close kin pairs found here fall within the calculated range of dispersal kernel spread for *Acropora* taxa on the Great Barrier Reefs calculated by Meziere *et al*. (2025). Larvae of many *Acropora* species experience precompetency periods of three to seven days before they are able to settle (Randall et al., 2024), and they may survive for 50 or even 100 days in the lab (Connolly & Baird, 2010; Miller et al., 2020; Nishikawa & Sakai, 2005). Dispersal models fit to genetic data suggest that dispersal times of more than 60 days are common in some *Acropora* species (Davies et al., 2015). Given that our study site exhibits mostly open coastlines, with surface current speeds that may exceed 0.5 meters per second during the seasonal monsoons, we can expect these competency times to result in long-distance larval transport (Catalano et al., 2024).

While three of these pairs were identified as possible clones by KING, this seems less plausible than close kin given known *Acropora* life history. *Acropora* do not produce asexual larvae and self-fertilize only rarely (Boutet & Schierwater, 2021; Willis et al., 1997). Clonal genotypes in other *Acropora* are most often found within a few meters of each other, as they are derived from physically fragmented adult corals (Howlett et al., 2024). However, some species of *Acropora* may be more prone to reproduction by fragmentation than others, and coral fragments have occasionally been documented to settle and grow after transportation by strong typhoons, such as those common in the Philippines (Anticamara & Tan, 2018; Pipithkul et al., 2021). It is possible though unlikely that these putative clone pairs result from fragmentation during strong storms and subsequent cross-island transportation.

Alternatively, coral polyps under acute stress occasionally detach themselves completely from the coral skeleton and form a “pseudo-planula” in a process known as “polyp bailout”. These detached polyps can later settle in a new location and grow into a new colony. Polyp bailout was originally documented in the 1940s (incidentally, in a species of *Acropora cf. tenuis*), but remains an understudied process in coral ecology (Schweinsberg et al., 2021). Whether the related individuals in our dataset represent true clones or first degree relatives, our finding of closely related individuals at widely spaced sites provides strong evidence of long-distance gene flow across a single generation.

### Algal symbionts structured more strongly by geography than host

*Cladocopium* taxonomy remains poorly resolved, with only a handful of formally recognized species out of potentially hundreds in nature (Butler et al., 2023; Davies et al., 2023). Some species of *Cladocopium* associate with specific coral hosts, especially but not exclusively hosts that transmit their symbionts vertically (Thornhill et al., 2014; Turnham et al., 2021). However, other named species of *Cladocopium* are generalists, found across thousands of kilometers and multiple hosts, including multiple species of *Acropora* (Butler et al., 2023). Extremely low FSTs between *Cladocopium* populations found in different coral taxa further imply that the taxonomic variation in our *Acropora* samples is not reflected in their algal symbionts. A study of co-occurring *Acropora cf. tenuis* cryptic taxa on the Great Barrier Reef also found a common pool of *Cladocopium* symbionts between cryptic taxa (Matias et al., 2023). Similarly in our study, we find a single genetic grouping of *Cladocopium* shared between all *Acropora* taxa. While ADMIXTURE resolved three ancestry groups, all groups were present in all *Acropora* taxa, showing structure with geography (latitude) rather than host. Our finding of weakly significant genetic differentiation between *Cladocopium* in different *Acropora* taxa could imply some symbiont selectivity by coral hosts, or it could result from common environmental factors structuring both host taxon occurrence and *Cladocopium* genotype. For example, we found that the *Acropora* taxa exhibited slight latitudinal differences in their distributions, and we found that latitude structured genetic variation in *Cladocopium* as well. This finding could result from neutral variation across the seascape, or it could represent adaptive genetic differences relating to differences in temperature along the latitudinal gradient.

Studies of *Acropora* taxa on other reefs have also found genetic variation in symbionts that is structured by environmental factors separately from coral host genomes. Genetic variation of *Cladocopium* found in *Acropora cf. humilis* in the Coral Sea depends largely on environmental variables such as thermal stress, light attenuation, and latitude, and only very slightly on host genotype (Marzonie et al., 2024). *Acropora hyacinthus* in Palau host mostly *Cladocopium*, and the genomes of those *Cladocopium* vary both between reefs and with depth (Armstrong et al., 2024). However, studies in other *Acropora* taxa, including *A. hyacinthus* and *A. digitifera* in Micronesia, have found host specialization between different *Cladocopium* lineages (Davies et al., 2020). It is worth noting that all of the *Acropora* taxa in our study are likely closely related, as evidenced by the relatively low FSTs between the groups. Other co-occurring *Acropora* species in the central Philippines may selectively host different lineages of *Cladocopium*, or even other symbiont genera. However, within this *A. cf. tenuis* species complex, genetic variation in algal symbionts appears to be more driven by geography and environment than host genetic variation. This result is consistent with study of *Acropora* cryptic taxa on the Great Barrier Reefs, where a common pool of *Cladocopium* found across hosts nonetheless exhibited significant genetic variation with distance to shore (Matias et al., 2023).

In our study, *Cladocopium* exhibited significant isolation by distance patterning in both one and two dimensions. Resampled IBD slopes were more consistently positive than those found in *Acropora*, even when controlling for sample size, for all of the taxa in one dimension and for three of four taxa in two dimensions. Our *Cladocopium* samples also exhibited spatial genetic variation at fine scales, as evidenced by the strong correlation between the second principle component of the algal genomes and depth. Partial Mantel tests also revealed that genetic distance between *Cladocopium* populations depended on geographic distance, when controlling for genetic distance between *Acropora* populations more strongly than the inverse. All of these lines of evidence indicate that within these *Acropora* populations, geography and environment structure symbiont genetic variation more strongly than does the genome of the host.

Despite these differences in genetic structure, the relative dispersal distances in the two taxa are difficult to measure directly. The slope of the relationship between geographic and genetic distance is inversely related to dispersal kernel spread, with steeper slopes implying shorter dispersal distances (Rousset, 1997). However, the slope is also inversely related to effective population size, with larger effective population sizes resulting in shallower slopes (Rousset, 1997). Without effective population size estimates for these taxa, dispersal cannot be estimated. In addition, differences in effective population size could explain some of the differences that we observed. For example, the weakly steeper slopes of some *Acropora* IBD relationships compared to the *Cladocopium* relationship could result from much higher effective population sizes in the latter. Microscopic *Cladocopium* can inhabit a variety of hosts and may also live freely in the water column or benthos (Bell & Quigley, 2025; Lim et al., 2019). The inverse relationship, however, is highly unlikely, namely that *Acropora* corals have substantially higher effective population sizes than *Cladocopium* in our study region. Therefore, the generally stronger isolation by distance patterns in *Cladocopium,* along with the observed patterns of landscape genetic variation, most likely indicates that *Acropora* disperse much further than their algal symbionts.

### Implications for gene flow and local adaptation

Corals are not the only marine animals that acquire photosymbionts from their settlement environment. Other examples include giant clams and jellyfish (Astorga et al., 2012; Butler et al., 2023; Fitt & Trench, 1981). In a very different marine environment, deep sea snails inhabiting hydrothermal vents also acquire their chemosynthetic bacterial partners horizontally (Breusing et al., 2022). Similarly to our finding that *Cladocopium* genotypes depend on geography and depth more strongly than host taxon, genetics of snail microsymbionts depend more strongly on site-specific environmental variables than on the genetics of their hosts. Environmental acquisition of shorter-dispersing symbionts can enable a far-dispersing host to acquire symbiont strains adapted to their specific local conditions (Breusing et al., 2022). Our results suggest the possibility of similar dynamics at play in *Acropora* in the Philippines. We found low genetic structuring in populations of four different *Acropora* taxa across a two hundred kilometer span on multiple islands, suggesting limited potential for genome-wide local adaptation. Conversely, we found genetic variation in *Cladocopium* associated with latitude and depth, both of which determine the environmental conditions to which corals are exposed.

Our results revealed differing patterns of spatial genetic variation in *Cladocopium* and their *Acropora* hosts, despite their close symbiotic relationship. The contrast between extensive within-species connectivity in *Acropora*, contrasted with the spatial geographic structuring in *Cladocopium,* creates evolutionary advantages for the holobiont. Long dispersal distances enable genetic connectivity between far-flung populations, but can limit local adaptation when parents and offspring settle in very different conditions (Kawecki & Ebert, 2004). This may be a particular problem in the marine realm, where species tend to exhibit longer dispersal distances than do terrestrial taxa (Kinlan & Gaines, 2003). Here we see *Acropora cf. tenuis* larvae traversing potentially hundreds of kilometers of latitudinal span, as well as tens of meters of depth gradient, and ultimately settling in markedly different environments than their parents. Acquiring symbionts from the settlement environment may allow far-dispersing marine taxa to take advantage of locally adapted symbiont genotypes.

Given these differences, simulations of gene flow and evolution in corals should consider modeling coral hosts and their algal symbionts as distinct yet interacting components within a single framework. The close ecological relationship between *Acropora* and their *Cladocopium* symbionts does not require parallel patterns of landscape genetics, as the taxa exhibit contrasting patterns of speciation, gene flow, and possibly adaptation. Continued study of close symbiont pairs, including corals, will be important for understanding how species interactions shape the co-evolution of dispersal.

## Acknowledgements

This study was made possible by funding from a National Defense Science and Engineering Graduate Fellowship (JTB), the National Science Foundation under award OCE-2443233 (LCM), and the Paul M. Angell Foundation under award CON-F23-28 (MLP and JTB). The authors thank Isabel M. Silva and Peter T. Raimondi for comments on an earlier version of this project.

## Data availability statement

Raw reads are downloadable from NCBI at https://www.ncbi.nlm.nih.gov/sra/PRJNA1445311. All code, metadata, and selected intermediates are available on Github at https://github.com/pinskylab/Atenuis_Philippines.

Github repository will be replaced with Zenodo link upon acceptance.

Metadata is additionally published on GEOME at https://n2t.net/ark:/21547/HDm2.

