## Supplemental tables and figures for "Contrasting patterns of seascape genetics in *Acropora cf. tenuis* and their symbiotic algae"

**S1. Genetic grouping and ADMIXTURE in *Acropora.***

**Supplemental Tables**

**Supplemental Table 1.** Samples per site. Sites on each island are numbered from north to south. A full list of coordinates per site can be found at https://github.com/pinskylab/Atenuis_Philippines/tree/main/metadat in the 'all_Atenuis_sites.csv' file. Each row here corresponds to a site along the coast of Cebu ("CEB") or Leyte ("LEY"). Four columns show the number of *Acropora* samples assigned to each taxon at each site, and the final column shows the number of samples with usable *Cladocopium* genomes at each site.

| Site | Taxon 1 count | Taxon 2 count | Taxon 3 count | Taxon 4 count | *Cladocopium* count |
|---|---|---|---|---|---|
| CEB01 | 2 | 3 | 3 | 5 | 8 |
| CEB02 | 2 | 0 | 1 | 0 | 1 |
| CEB03 | 0 | 0 | 0 | 1 | 2 |
| CEB04 | 1 | 0 | 4 | 1 | 9 |
| CEB05 | 4 | 1 | 1 | 0 | 10 |
| CEB06 | 5 | 1 | 0 | 5 | 11 |
| CEB07 | 3 | 5 | 0 | 3 | 5 |
| CEB08 | 4 | 5 | 2 | 6 | 9 |
| CEB09 | 3 | 4 | 1 | 2 | 8 |
| CEB10 | 7 | 2 | 0 | 0 | 5 |
| CEB11 | 3 | 5 | 1 | 0 | 1 |
| CEB12 | 4 | 1 | 0 | 0 | 7 |
| CEB13 | 1 | 0 | 0 | 3 | 8 |
| CEB14 | 4 | 0 | 0 | 2 | 2 |
| CEB15 | 0 | 0 | 1 | 1 | 1 |
| CEB16 | 4 | 0 | 2 | 3 | 10 |
| CEB18 | 1 | 0 | 0 | 2 | 1 |
| CEB19 | 1 | 2 | 2 | 2 | 1 |
| CEB20 | 0 | 1 | 2 | 1 | 11 |
| CEB21 | 1 | 0 | 0 | 0 | 0 |
| CEB22 | 7 | 1 | 1 | 1 | 6 |
| CEB23 | 3 | 1 | 0 | 1 | 4 |
| CEB24 | 1 | 3 | 1 | 0 | 11 |

| | | | | | |
|---|---|---|---|---|---|
| CEB25 | 4 | 0 | 4 | 0 | 6 |
| CEB26 | 6 | 1 | 10 | 1 | 2 |
| CEB27 | 3 | 1 | 1 | 1 | 8 |
| CEB29 | 3 | 0 | 1 | 1 | 14 |
| CEB30 | 0 | 0 | 0 | 0 | 3 |
| CEB32 | 5 | 0 | 0 | 0 | 11 |
| CEB33 | 1 | 7 | 0 | 0 | 7 |
| CEB35 | 2 | 5 | 3 | 0 | 9 |
| LEY01 | 1 | 0 | 0 | 0 | 11 |
| LEY02 | 12 | 0 | 0 | 0 | 14 |
| LEY04 | 2 | 3 | 0 | 0 | 16 |
| LEY05 | 1 | 1 | 2 | 1 | 12 |
| LEY06 | 1 | 0 | 0 | 1 | 12 |
| LEY07 | 5 | 0 | 1 | 0 | 11 |
| LEY09 | 0 | 0 | 2 | 0 | 8 |
| LEY10 | 0 | 5 | 5 | 0 | 11 |
| LEY11 | 1 | 0 | 1 | 1 | 4 |
| LEY13 | 3 | 4 | 2 | 2 | 4 |
| LEY14 | 0 | 4 | 1 | 4 | 4 |
| LEY15 | 5 | 3 | 2 | 2 | 12 |

**Supplemental Table 2.** Genetic clustering in *Acropora* with Discriminant Analysis of Principal Components and ADMIXTURE. The four genetic cluster solution minimizes the BIC for the DAPC, indicating the best model fit. Likewise, the lowest cross-validation error for the ADMIXTURE grouping occurs with four genetic ancestry groups.

| Number of groups (K=) | BIC from DAPC clustering | ADMIXTURE cross-validation error |
|---|---|---|
| 1 | 1573.9 | 0.38265 |
| 2 | 1553.9 | 0.34834 |
| 3 | 1542.8 | 0.33183 |
| 4 | **1535.2** | **0.32198** |
| 5 | 1535.3 | 0.32660 |
| 6 | 1539.2 | 0.32744 |
| 7 | 1543.5 | 0.33064 |
| 8 | 1546.8 | 0.33429 |
| 9 | 1550.9 | 0.33827 |
| 10 | 1554.9 | 0.34398 |
| 11 | 1558.8 | 0.34672 |
| 12 | 1562.6 | 0.35126 |
| 13 | 1566.5 | 0.35800 |
| 14 | 1570.7 | 0.36272 |
| 15 | 1574.6 | 0.36438 |
| 16 | 1578.9 | 0.36977 |

**Supplemental Table 3.** Individual samples with the highest number of symbiont reads mapping to *Durusdinium*, rather than *Cladocopium* or *Symbiodium*. For three individuals, the total number of reads mapped was negligible (<1000), indicating poor sequencing, and two more individuals showed slightly higher but still low (<50,000) numbers of mapped reads. Two samples (CEB01_T14 and CEB20_T08) presented a high (>150,000) number of reads, with a substantial majority (>80%) mapping to *D. trenchii* rather than other symbiont genomes (bold). The *Acropora* taxon column shows the taxonomic group to which the host was assigned. Four of the hosts lacked sufficient SNPs to be confidently assigned to any taxon. The depth column shows the collection depth for the host coral in feet.

| Individual ID | Number of reads mapped to *Durusdinium* | % symbiont reads mapped to *Durusdinium* | *Acropora* taxon | Depth |
|---|---|---|---|---|
| **CEB01_T14** | **455033** | **92%** | **1** | **NA** |
| CEB04_T05 | 64 | 49% | NA | NA |
| CEB11_T08 | 28873 | 87% | NA | NA |
| CEB16_T04 | 260 | 59% | NA | NA |
| CEB18_T20 | 194 | 51% | NA | NA |
| **CEB20_T08** | **172902** | **82%** | **4** | **NA** |
| LEY02_T02 | 31833 | 58% | 1 | 13 ft. |

**Supplemental Table 4.** Individual samples with the highest number of symbiont reads mapping to *Symbiodium*, rather than *Cladocopium* or *Durusdinium*. One individual coral sample had a majority of symbiont reads mapping to the *S. kawagutii* genome.

| Individual ID | Number of reads mapped to *Symbiodinium* | % symbiont reads mapped to *Symbiodinium* | *Acropora* taxon | Depth |
|---|---|---|---|---|
| CEB11_T11 | 22467 | 64% | 1 | NA |

**Supplemental Table 3.** Genetic clustering in *Cladocopium* with Discriminant Analysis of Principal Components and ADMIXTURE. A single genetic cluster minimizes the BIC for the DAPC, indicating the best model fit. The lowest cross-validation error for the ADMIXTURE grouping occurs with three genetic ancestry groups.

| Number of groups (K=) | BIC from DFA clustering | ADMIXTURE cross-validation error |
|---|---|---|
| 1 | **1520.3** | 0.45678 |
| 2 | 1520.7 | 0.45692 |
| 3 | 1521.7 | **0.45349** |
| 4 | 1524.3 | 0.45647 |
| 5 | 1527.0 | 0.45629 |
| 6 | 1530.0 | 0.46088 |
| 7 | 1533.5 | 0.46570 |
| 8 | 1537.0 | 0.46804 |
| 9 | 1540.5 | 0.46868 |
| 10 | 1544.4 | 0.47380 |
| 11 | 1548.1 | 0.47189 |
| 12 | 1552.0 | 0.47404 |
| 13 | 1556.0 | 0.47852 |
| 14 | 1560.0 | 0.48440 |
| 15 | 1563.8 | 0.48839 |
| 16 | 1567.8 | 0.48202 |

**Supplemental Table 6.** One dimensional IBD slopes and Mantel tests along the coast of Cebu. P-values marked with ** are highly significant (p<0.01).

| **Taxon** | **Slope** | **Standard error** | **Mantel test p-value** |
|---|---|---|---|
| *Acropora* Taxon 1 | $-8.4 \times 10^{-9}$ | $1.2 \times 10^{-8}$ | 0.683 |
| *Acropora* Taxon 2 | $-2.2 \times 10^{-8}$ | $1.5 \times 10^{-8}$ | 0.681 |
| *Acropora* Taxon 3 | $4.9 \times 10^{-8}$ | $8.5 \times 10^{-8}$ | 0.273 |
| *Acropora* Taxon 4 | $3.1 \times 10^{-7}$ | $2.6 \times 10^{-7}$ | 0.160 |
| *Cladocopium* | 9.2e-08 | $1.8 \times 10^{-8}$ | 0.001** |

**Supplemental Table 7.** Two dimensional IBD slopes and Mantel tests from across the entire study area, including Cebu, Leyte, and the Camotes islands. P-values marked with * are marginally significant (p<0.1) and p-values marked with ** are highly significant (p<0.01).

| **Taxon** | **Slope** | **Standard error** | **Mantel test p-value** |
|---|---|---|---|
| *Acropora* Taxon 1 | -0.00031 | 0.00053 | 0.679 |
| *Acropora* Taxon 2 | -0.00054 | 0.00074 | 0.710 |
| *Acropora* Taxon 3 | 0.0056 | 0.0043 | 0.064* |
| *Acropora* Taxon 4 | 0.017 | 0.014 | 0.174 |
| *Cladocopium* | 0.0055 | 0.0014 | 0.004** |

**Supplemental Table 8.** Putative parent offspring pairs.

| **Individual 1** | **Individual 2** | **Taxon** | **Opposite homozygotes** | **SNPs in both** | **LLR** | **Geographic distance** |
|---|---|---|---|---|---|---|
| **LEY07_T02** | **CEB35_T08** | **1** | **0** | **1245** | **16.12** | **58 km** |
| LEY11_T15 | LEY04_T15 | 1 | 0 | 2284 | 13.88 | 45 km |
| **LEY14_T09** | **CEB07_T14** | **2** | **1** | **1649** | **26.00** | **142 km** |
| CEB26_T22 | CEB05_T12 | 3 | 0 | 1200 | 14.94 | 174 km |
| **CEB01_T12** | **CEB23_T16** | **4** | **0** | **1075** | **5.46** | **181 km** |
| **CEB01_T11** | **CEB23_T16** | **4** | **0** | **1074** | **4.99** | **181 km** |
| **CEB01_T11** | **CEB01_T12** | **4** | **0** | **1159** | **4.79** | **0 km** |

**Supplemental Figures**

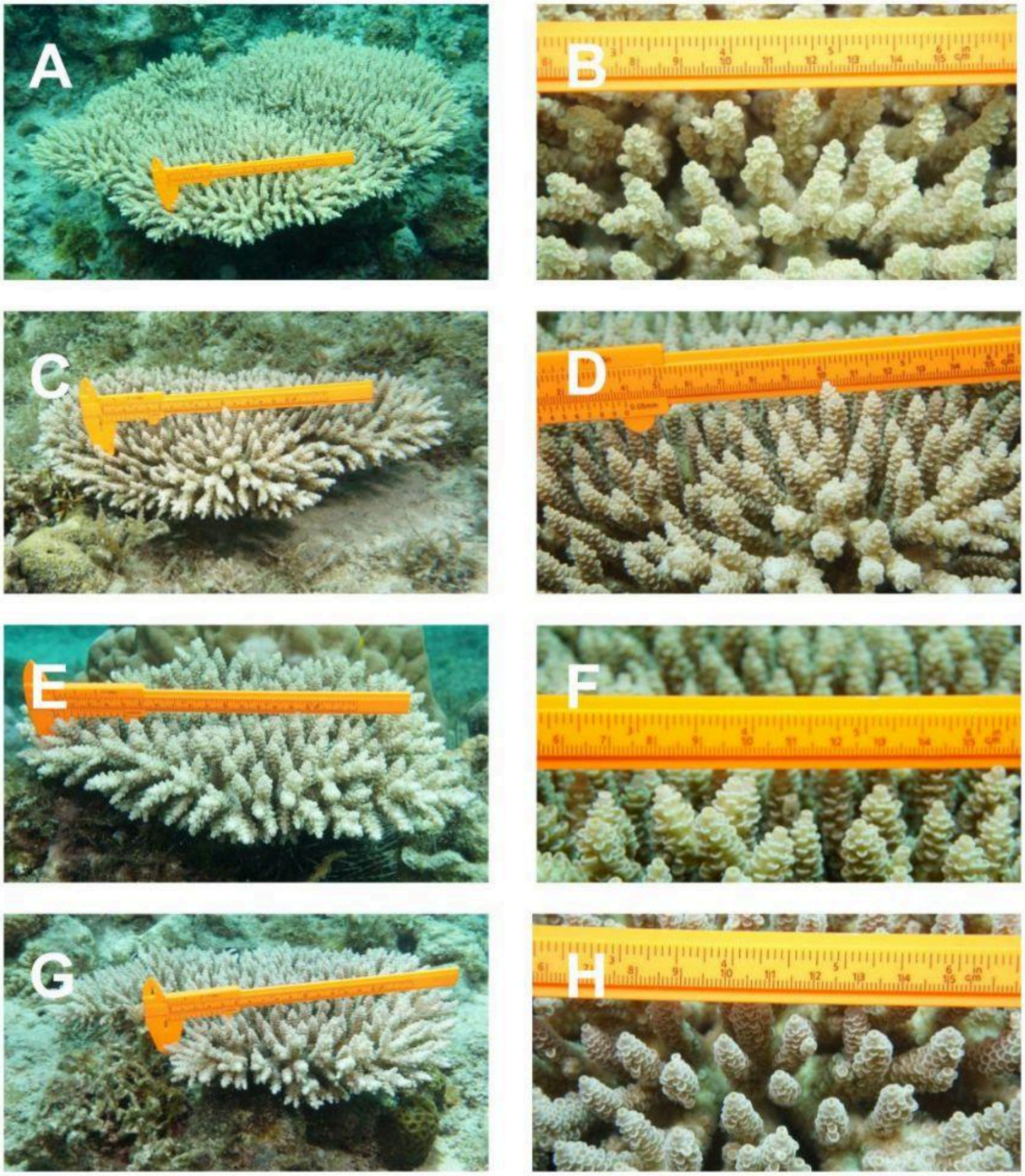


**Supplemental Fig. 1.** Cryptic taxa in *Acropora cf. tenuis* are morphologically similar. Each row shows one representative sample from one cryptic taxon. The top row (**A** and **B**) shows Taxon 1, the second row shows Taxon 2, the third row shows Taxon 3, and the fourth row shows Taxon 4. The left column (**A**, **C**, **E, G)** shows overall growth form and the right column (**B**, **D**, **F**, **H**) shows close ups of corallite structure.

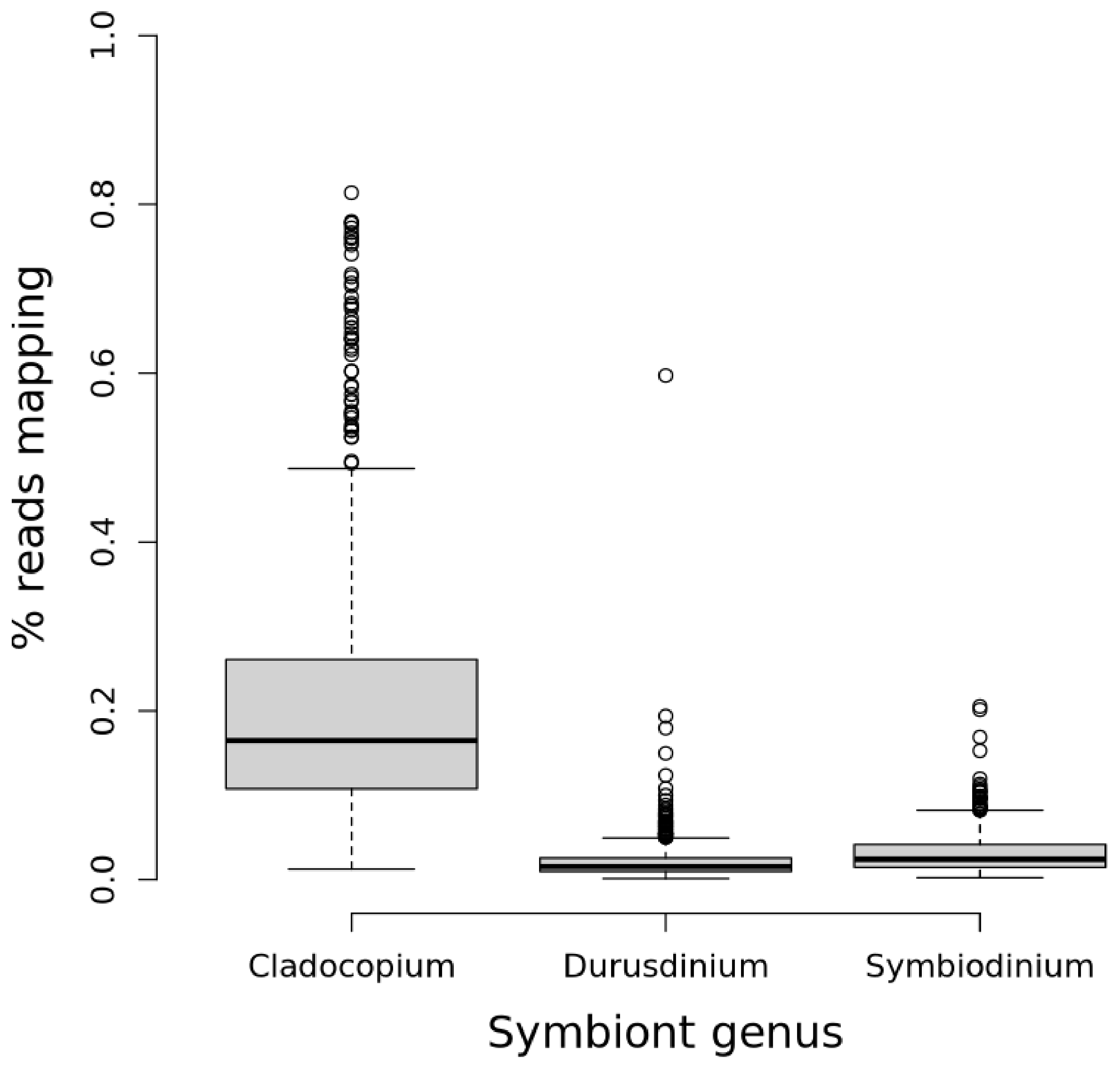


**Supplemental Fig 2.** Across samples, more reads mapped to the genome from the *Cladocopium* genus than genomes from the genera *Durusdinium* or *Symbiodinium*. A median of 14,424.5 reads, or 15% of the total, successfully mapped to the *Cladocopium goreaui* genome, 2771 reads (2.4% of total) mapped to the *Symbiodinium kawagutii* genome, and 1447 reads (1.5% of total) mapped to the *Durusdinium trenchii* genome.

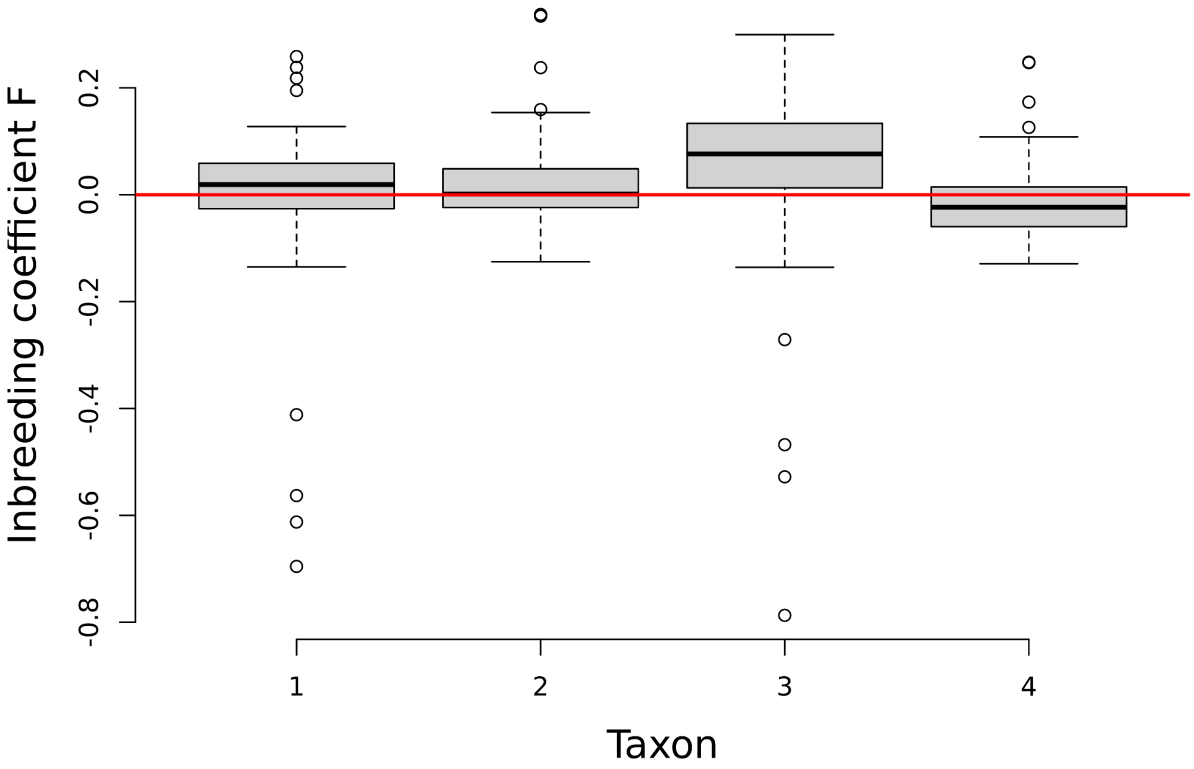


**Supplemental Fig 3.** Cryptic taxa in *Acropora* show different mean inbreeding coefficients (Kruskall-Wallis test p=3.1 x $10^{-6}$, n=295). The horizontal red line shows zero inbreeding (expected heterozygosity).
